## Supplementary Figures for "Cell-attribute aware community detection improves differential abundance testing from single-cell RNA-Seq data"

### SUPPLEMENTARY INFORMATION

#### This document contains all Supplementary Figures and their legends.

#### Supplementary Figures

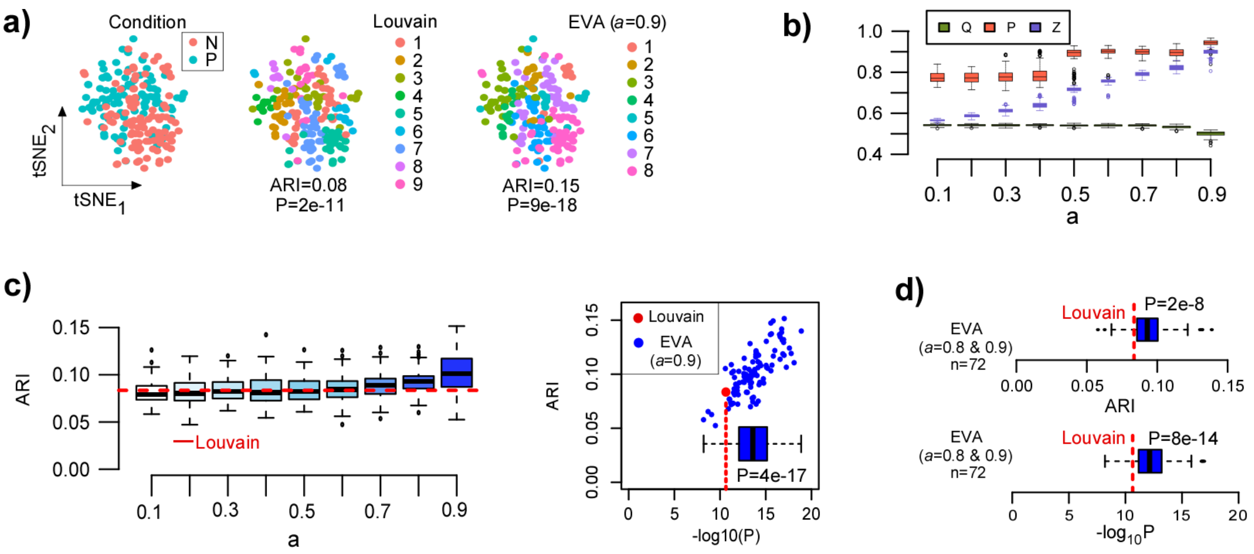


**SI fig.S1: Testing of the EVA R-implementation on simulated data: a)** tSNE visualization of a simulated scRNA-Seq dataset consisting of 200 cardiomyocyte cells from one mouse with half of the cells representing a perturbation (P) state. N=normal. In middle and right panels, cells are annotated according to the clusters inferred with Louvain and EVA (a=0.9) on the nearest-neighbor cell-cell graph. For EVA the result of only one run is shown. The Adjusted Rand Index (ARI) of clusters against condition is given, as well as the Chi-Square statistic P-value of association between clusters and condition. **b)** Boxplots displaying the modularity (Q), the purity (P) and generalized modularity (Z) as a function of purity index parameter a for EVA. Each boxplot contains the values of 100 distinct EVA runs. **c)** Boxplots displaying the ARI as a function of purity index parameter a. Each boxplot contains the values of 100 distinct EVA runs. Red dashed line indicates the ARI derived using Louvain. Right panel represents a scatterplot of the ARI (y-axis) against the statistical significance of Chi-Square statistic P-values (x-axis, -log10P). The inlet boxplot summarizes the statistical significance values and P-value is from a one-sided Wilcoxon rank sum test comparing these values from EVA against the Louvain null value. **d)** Boxplots comparing the ARI and statistical significance values of EVA to the Louvain values, when restricting to 72 EVA runs (with a=0.8 or a=0.9) for which the number of inferred communities equals the number of communities inferred with Louvain (n=9). P-values are from one-sided Wilcoxon rank sum tests comparing the 72 EVA derived values to the Louvain null.

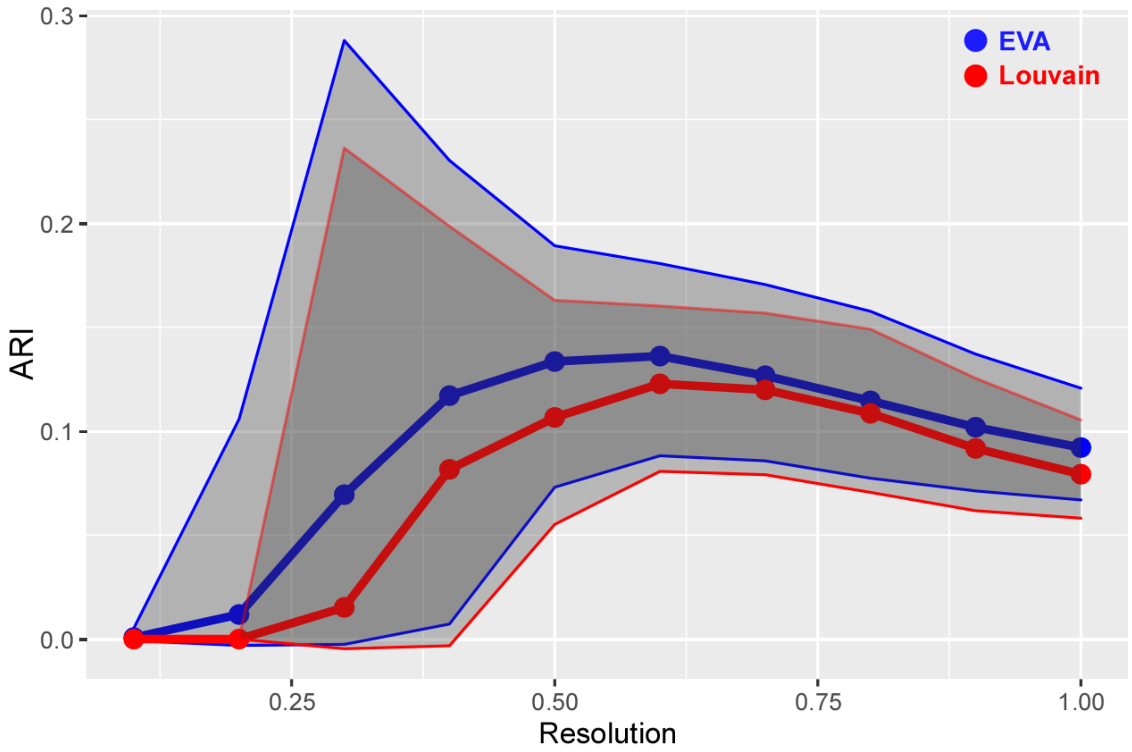


**SI fig.S2: Comparison of EVA and Louvain in relation to resolution parameter:** Using the same simulated data as in SI fig.S1, a plot of the adjusted Rand Index (ARI) of EVA and the Louvain algorithm against the resolution parameter. Data points represent averages over 1000 distinct instances/runs of each algorithm. The lines define 95% CI envelopes.

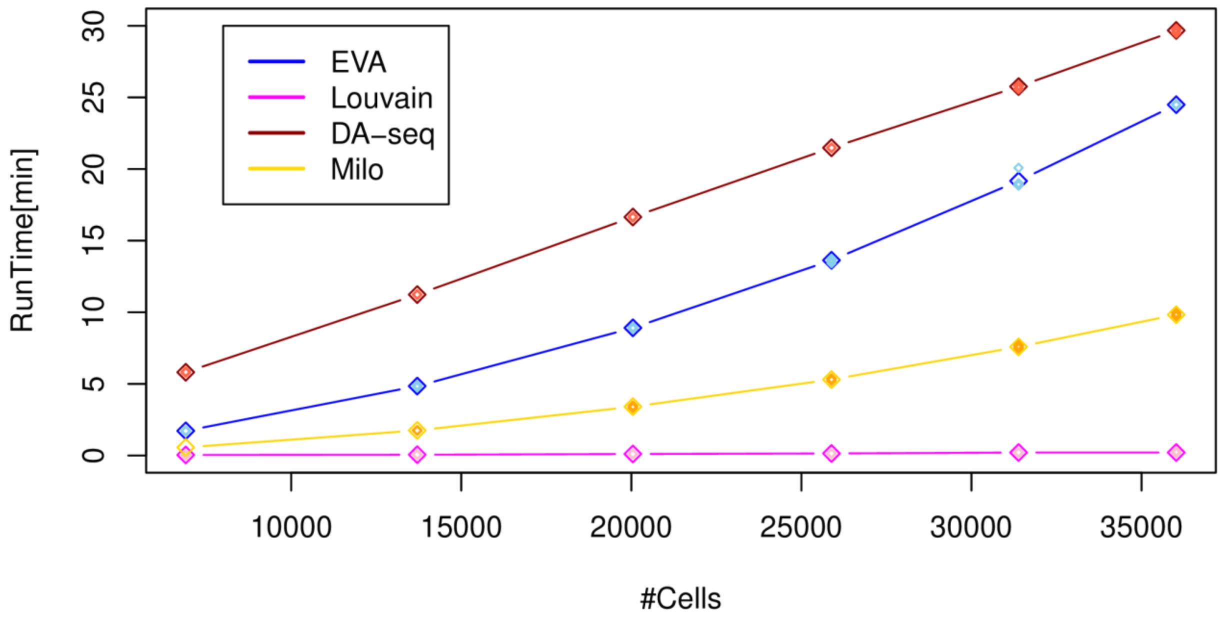


**SI fig.S3: Runtime comparison of EVA.** Runtimes for one run in minutes (y-axis) vs. the number of cells (#Cells, x-axis), as estimated using the enterocyte snRNA-Seq dataset encompassing a maximum of 103913 cells, and for 4 different methods as shown. In the case of EVA and Louvain, the number of cells defines the size of the cell-cell *k*-nn graph, where *k*=50 was kept fixed throughout. In the case of DA-seq, because it requires a binary phenotype, we ran 3 separate comparisons for DA (normal+unaffected vs polyp, polyp vs adenoma, and normal+unaffected vs adenoma). Of note, for EVA and Louvain the timing measures the time to find communities given the input graph. To make the comparison objective and fair, the timing for DAseq was taken as the time to run the functions getDAcells and getDAregion, which does the inference of enriched regions, analogous to finding the enriched communities in the case of EVA. Likewise, for Milo the timing was measured from the construction of neighborhoods (makeNhoods function), ending with the neighborhood annotation (annotateNhoods function) which identifies enriched neighborhoods. For detailed parameter choices, see Methods section. All runtimes were obtained on a Dell Precision Workstation with an Intel Xeon(R) CPU E3-1575M v5 @3GHz and 64 GB RAM. Although workstation comes with 8 processing cores, runtimes were obtained without parallelization.

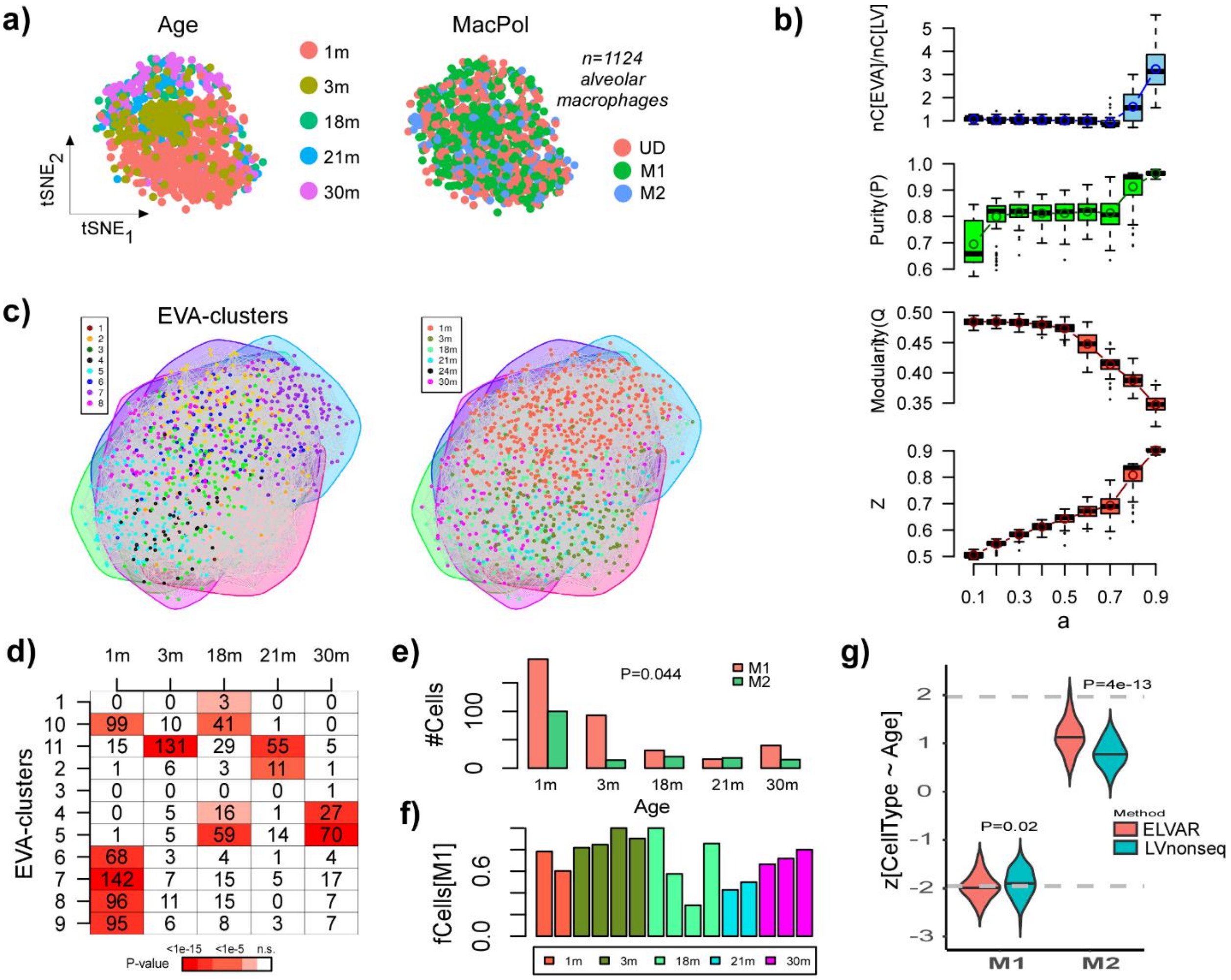


**SI fig.S4: ELVAR predicts age-related decrease in M1 polarization. a)** tSNE visualization of 1124 lung alveolar macrophages with cells annotated by age-group (left) and polarization subtype (right). M1/2=type-1/2 polarization, UD=undetermined. **b)** Top panel: boxplots display the number of communities inferred using EVA against the purity parameter $a$ , normalized relative to the number of communities inferred with the Louvain algorithm ($a=0$). Each boxplot represents the distribution over 100 different runs. Top middle panel: As top-panel but now with y-axis displaying the purity of the clusterings. Lower middle panel: As top-panel but now with y-axis displaying the modularity of the clusterings. Lower panel: as top-panel but now with y-axis displaying the objective function to be optimized. **c)** Left panel: Cell-cell nearest neighbor graph inferred using Seurat, with cell colors indicating the inferred EVA communities from one typical run. Right panel: as left-panel, but with cells now colored by age-group. **d)** Matrix entries give the number of cells per EVA-cluster and age-group, with color indicating the P-value of enrichment, for one particular run. For a given age-group, only cells from enriched clusters are taken forward using a Bonferroni-adjusted threshold (typically around 0.001). **e)** Barplot displaying the number M1 and M2 cells per age-group only using enriched clusters from d). P-value is from a two-tailed

Fisher-test. **f)** Barplots display the fraction of M1 cells for each age-group, with bars of the same color indicating different mouse replicates. **g)** Violin plots displaying the z-statistics (100 runs) derived from the negative binomial regression for the case of M1 and M2 cell-type fractions, with dashed horizontal lines indicating the P=0.05 significance level. Violin plots compare the z-statistics derived from ELVAR with those derived from an analogous algorithm that uses the non-sequential Louvain algorithm in place of EVA. P-values derive from a one-tailed Wilcoxon rank sum test comparing these two z-statistic distributions.

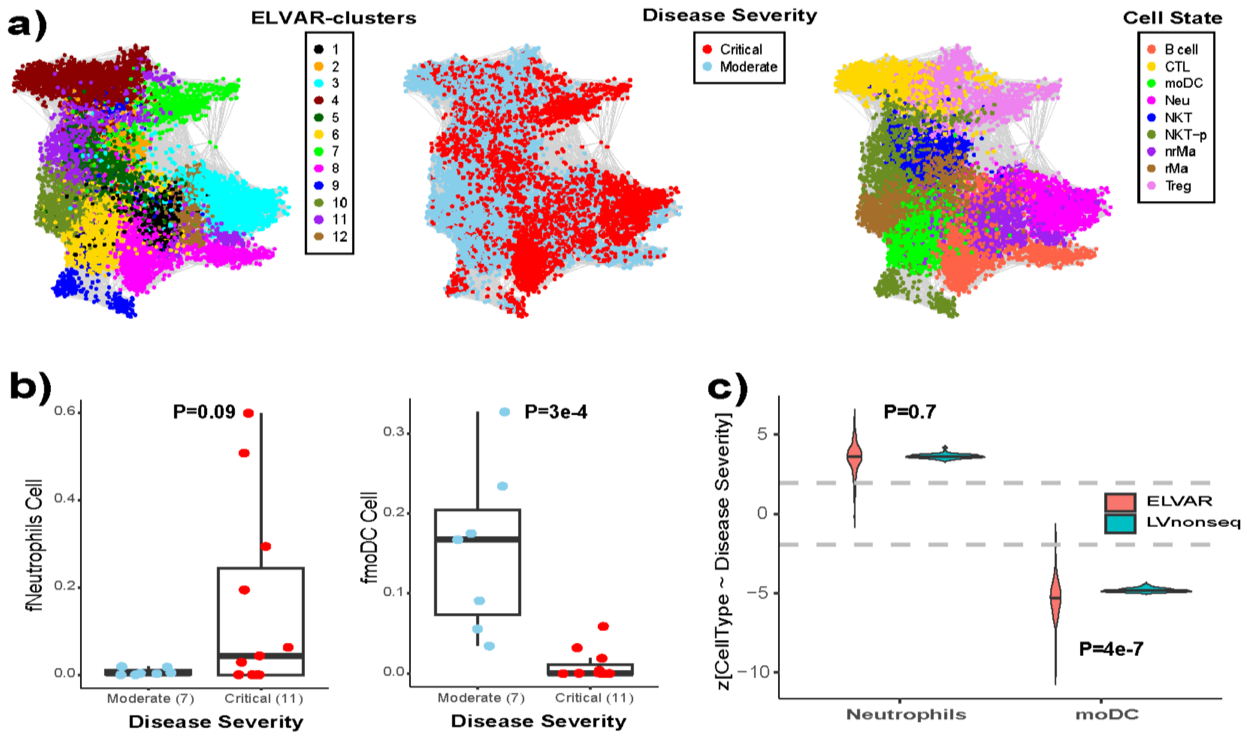


**SI fig.S5: ELVAR detects DA of immune cell-types with Covid-19 disease severity. a)** Left panel: The cell-cell similarity graph inferred using Seurat on scRNA-Seq data of immune cells annotated by community membership, as inferred using ELVAR. Middle and Right panels depict the same graph but with cells annotated by Covid-19 disease state (Moderate, Critical) and cell-type. Data are shown for one representative ELVAR run. **b)** Left panel: Boxplot displaying the neutrophil fraction as a function of disease severity, considering only cells that are part of significantly enriched ELVAR-clusters. P-value derives from a t-test. Right panel depicts the same but for monocyte-derived dendritic cells (moDCs). **c)** Violin plots display the z-statistics of the negative binomial regression of cell-count against disease severity for ELVAR and the analogous algorithm that uses non-sequential Louvain in place of EVA, with horizontal dashed lines indicating the P=0.05 level of statistical significance. P-values shown derive from a one-tailed Wilcoxon rank sum test comparing the ELVAR distribution to the Louvain-benchmark.

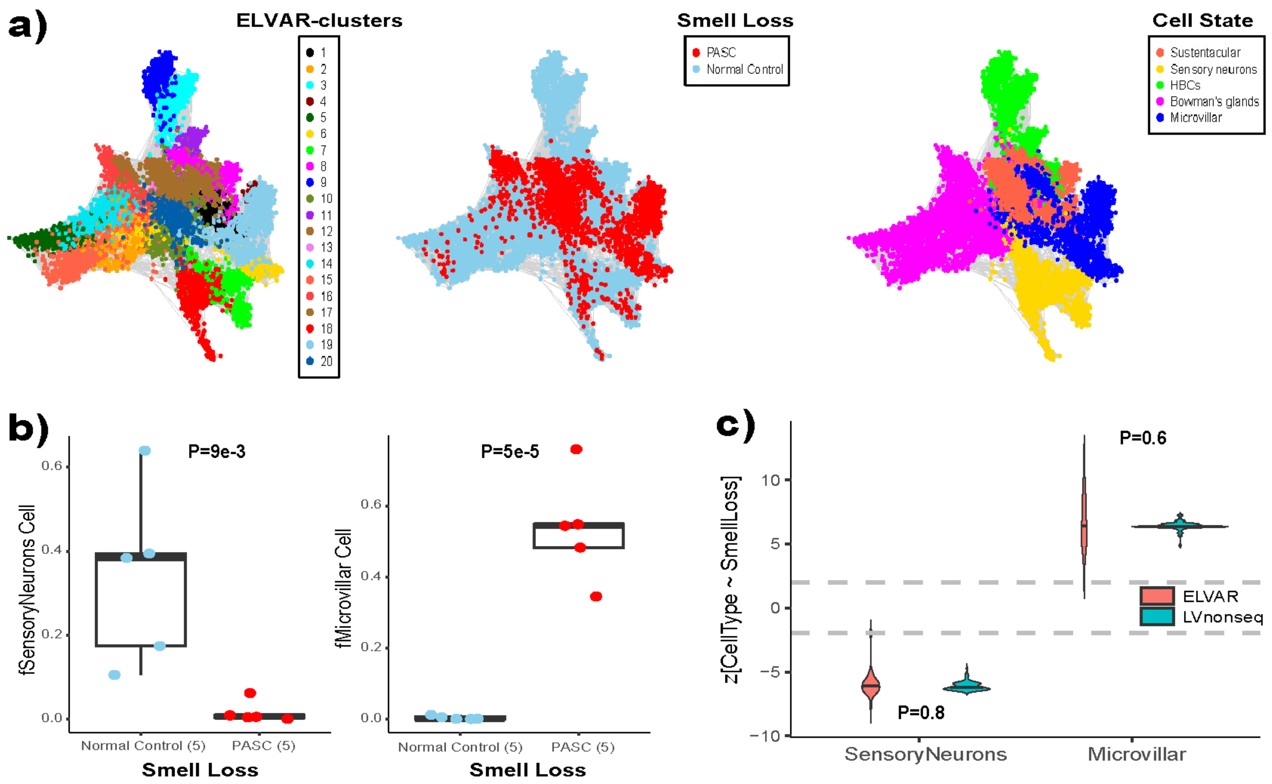


**SI fig.S6: ELVAR detects DA of cell-types associated with smell-loss in Covid-19 patients. a)** Left panel: The cell-cell similarity graph inferred using Seurat on scRNA-Seq data of olfactory epithelial cells annotated by community membership, as inferred using ELVAR. Middle and Right panels depict the same graph but with cells annotated by smell-loss Covid-19 phenotype (Normal: no smell loss, PASC: experienced smell loss) and cell-type. Data are shown for one representative ELVAR run. **b)** Left panel: Boxplot displaying the sensory neuron fraction as a function of smell-loss phenotype, considering only cells that are part of significantly enriched ELVAR-clusters. P-value derives from a t-test. Right panel depicts the same but for the microvillar cell fraction. **c)** Violin plots display the z-statistics of the negative binomial regression of cell-count against disease severity for ELVAR and the analogous algorithm that uses non-sequential Louvain in place of EVA, with horizontal dashed lines indicating the P=0.05 level of statistical significance. P-values shown derive from a one-tailed Wilcoxon rank sum test comparing the ELVAR distribution to the Louvain-benchmark.

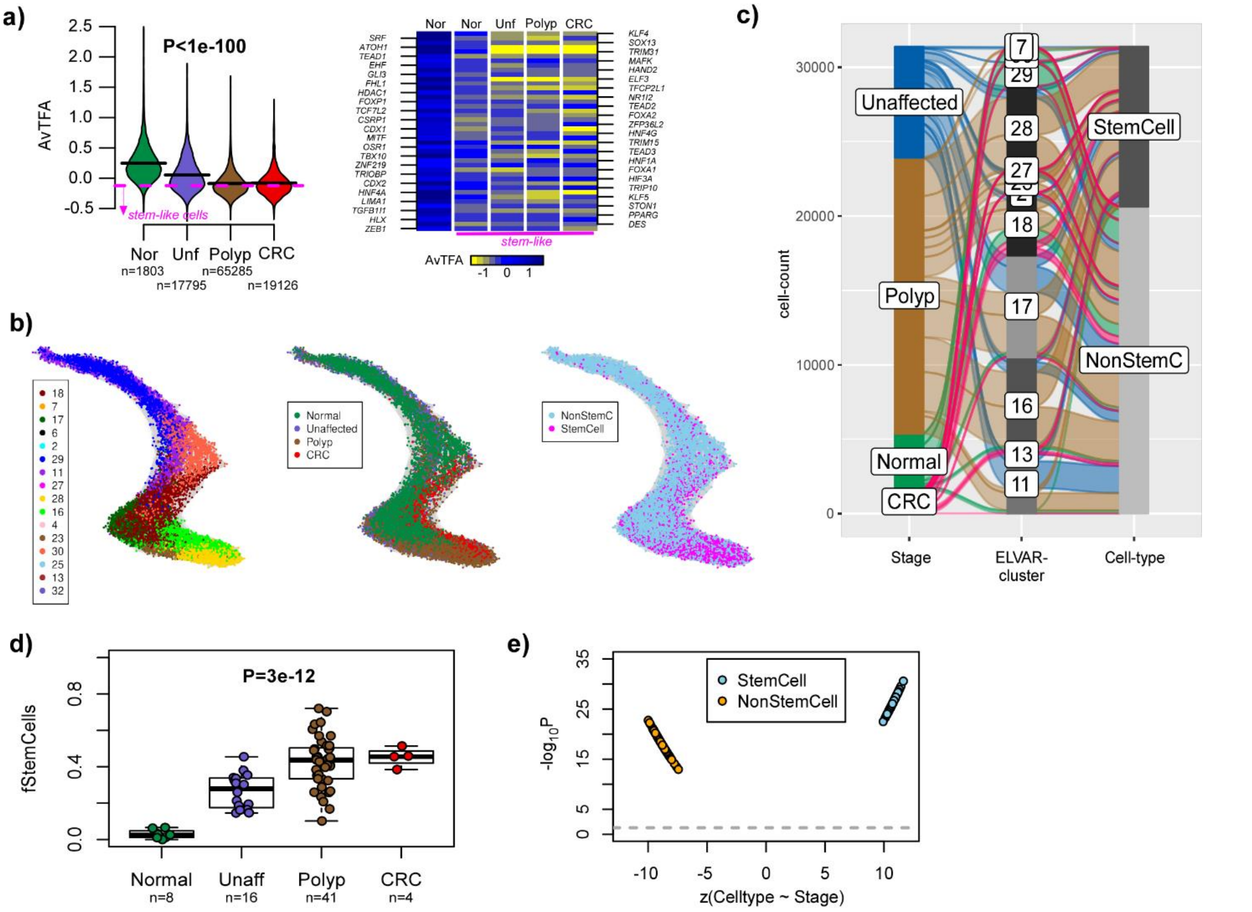


**SI fig.S7: ELVAR validation of increased stem cell fraction in polyps. a)** Left panel: Violin plots comparing the average differentiation activity (AvTFA), as estimated over 56 colon-specific TFs, across the four disease stages (Nor=normal, Unf=unaffected FAP cases, Polyp=predominantly FAP cases with polyps, CRC=colorectal adenocarcinoma (predominantly non-FAP)). P-value is two-sided from a linear regression. Pink dashed line indicates the 95% lower quantile of AvTFA values. Number of cells in each disease stage is indicated. Right panel: Heatmap of TFA values for a subset of 44 colon-specific TFs that display lower TFA in unaffected + polyp-carrying FAP cases compared to normal, with TFA values averaged over all normal cells, normal stem-like cells, unaffected stem-like cells, polyp stem-like cells and CRC stem-like cells. The stem-like cells were defined as in a). **b)** Cell-cell nearest neighbor graph (k=50) with cells annotated by inferred EVA/ELVAR community, disease stage and cell-state (stem-like vs non-stem cell). Data is shown for one EVA run. **c)** Alluvial plot for the same EVA run, displaying the composition of communities according to disease stage and cell-state. **d)** Boxplot comparing the fraction of stem-like cells in each disease stage, with the fraction computed for each independent sample using only cells from enriched communities. P-value is two-sided from a linear regression. **e)** Corresponding negative binomial regression analysis, displaying the level of statistical significance (y-axis) to the z-statistics of cell counts vs disease stage (x-axis). For each cell-state, there are 100 values representing 100 distinct ELVAR runs. Grey dashed line indicates the -log10(0.05) significance level.

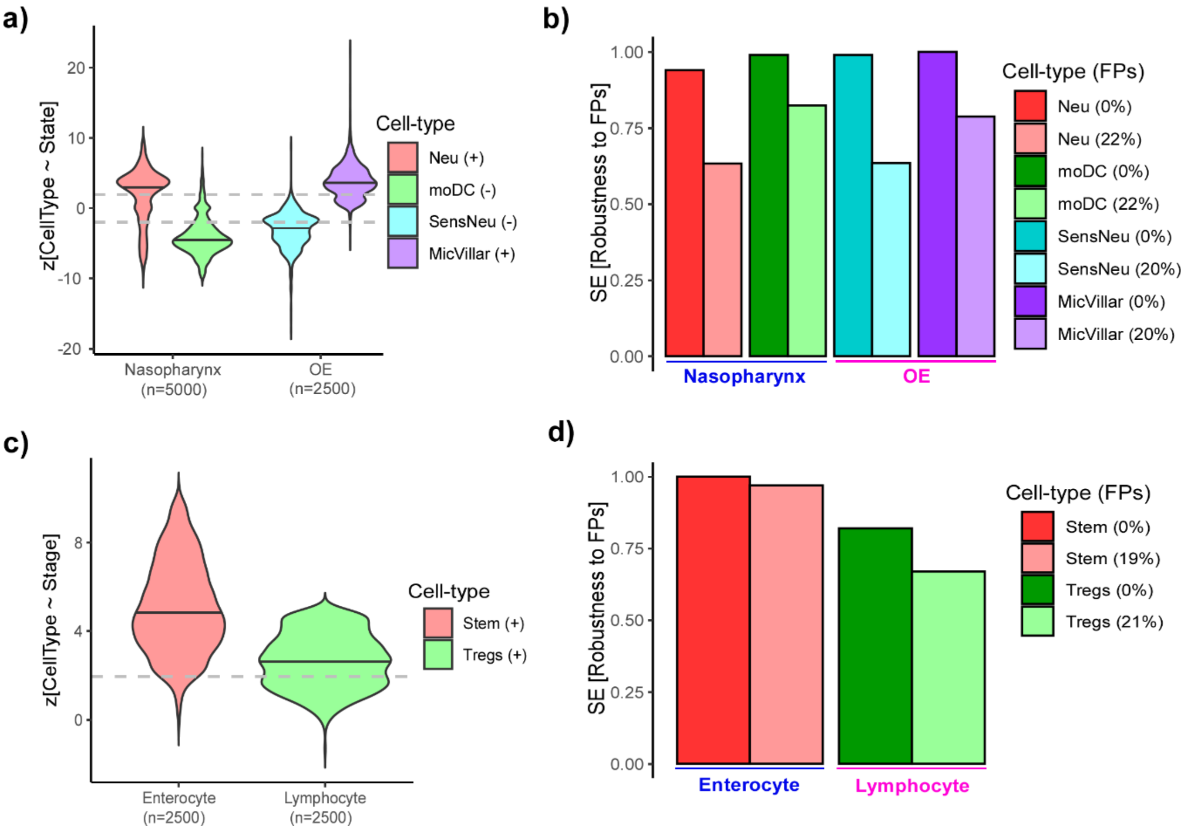


**SI fig.S8: Robustness of ELVAR to errors in clustering attribute. a)** Violin plots of the negative Binomial regression Wald z-statistics of association between cell-type counts and Covid-19 disease state (y-axis) obtained after introducing false positives (FP) in the clustering attribute (disease state). For the two Covid-19 datasets (nasopharynx & olfactory epithelium-OE) the fraction of false positives was 22% and 20%, respectively. In the case of OE, we had 5 controls and 5 cases, so we generated 25 perturbations in which 1 control and case were flipped to generate 20% FPs. For each such perturbation we performed a 100 ELVAR runs, so a total of 2500 runs. Similarly, for nasopharynx, we randomly flipped 2 cases and controls (corresponding to 22% FPs) a total of 50 times, each time performing 100 ELVAR runs, so a total of 5000 runs. For each tissue-type, we display the two main cell-types for which a DA-association had been found without FPs. The sign after the cell-type abbreviation indicates the directionality of the association observed without FPs. Grey dashed line corresponds to the line +/-1.96 (P=0.05). **b)** Barplot comparing the sensitivities of detecting a significant association (Wald z-statistic with P<0.05) with the same directionality as in the unperturbed case, as computed over the total number of runs. For comparison, we also display the sensitivities when no FPs are present. **c-d)** As a-b), but for the colon-polyp dataset focusing on enterocyte stem-cell and T-regulatory cells.
